# Combining molecular dynamics simulations and X-ray scattering techniques for the accurate treatment of protonation degree and packing of ionizable lipids in monolayers

**DOI:** 10.1101/2023.08.10.552652

**Authors:** Miriam Grava, Mohd Ibrahim, Akhil Sudarsan, Julio Pusterla, Julian Philipp, Joachim O. Rädler, Nadine Schwierz, Emanuel Schneck

## Abstract

The pH-dependent change in protonation of ionizable lipids is crucial for the success of lipid-based nanoparticles as mRNA delivery systems. Despite their widespread application in vaccines, the structural changes upon acidification are not well understood. Molecular dynamics simulations support structure prediction but require an a-priori knowledge of the lipid packing and protonation degree. The presetting of the protonation degree is a challenging task in the case of ionizable lipids since it depends on pH and on the local lipid environment and often lacks experimental validation. Here, we introduce a methodology of combining all-atom molecular dynamics simulations with experimental total-reflection X-ray fluorescence and scattering measurements for the ionizable lipid Dlin-MC3-DMA (MC3) in POPC monolayers. This joint approach allows us to simultaneously determine the lipid packing and the protonation degree of MC3. The consistent parameterization is expected to be useful for further predictive modeling of the action of MC3-based lipid nanoparticles.

## I. INTRODUCTION

The pH-dependent change in protonation of ionizable lipids is fundamental for endosomal release and hence the success of lipid nano-particles (LNPs) as mRNA delivery systems^1^. Ionizable lipids are already successfully used in FDA-approved drugs for the treatment of amyloidosis and in Moderna’s vaccines against SARS-CoV-2^2,3^. Hence, the design of ionizable lipids with high transfection efficiencies is expected to play a key role in modern RNA therapeutics. Despite their importance, the microscopic structural changes upon acidification, preceding the endosomal escape of RNA, have not been resolved so far. Molecular dynamics (MD) simulations are particularly suited to provide atomistic insights, however, they require an a-priori knowledge of the lipid packing and the protonation degree. The presetting of the protonation degree is not always a simple task since it depends sensitively on the environment. Three competing interactions have to be considered: (i) conformational rearrangements of the lipids, (ii) partial dehydration of the ionizable group, and (iii) interactions with polar and charged lipids or ions. Due to these contributions, it is not surprising that ionizable lipids can have significant pK_a_ shifts. For example, the pK_a_ value of Dlin-MC3-DMA (MC3) was found to be shifted from about 9.4 at infinite dilution^4^ to 6.4 inside a lipid nano-particle^5^.

Several theoretical models are available for pK_a_ predictions. Popular approaches are based on solving the linearized Poisson-Boltzmann equation^6^ or generalized Born models^7^. Such simple models work well for simple systems but fail in more complicated cases where conformational rearrangements become important. MD simulations include these conformational rearrangements and, when combined with free energy methods such as free energy perturbation^8^ or thermodynamic integration^9^, yield the free energy of deprotonation and hence the pK_a_. Using implicit or explicit solvent, free energy simulations have been successfully applied to calculate pK_a_ shifts in proteins^10,11^. In addition, constant-pH MD simulations have been developed to overcome limitations of fixed protonation states in conventional MD simulations^12,13^. However, these methods are computationally expensive and their accuracy depends on the underlying force fields and the convergence of the sampling of all degrees of freedom. In particular in lipid systems, which show pH dependent structural phase transitions^14^, such convergence is difficult to achieve. In addition, the correct packing of the lipids is crucial. To correctly reproduce the lateral structure, the force fields of all components including ionizable lipids, uncharged lipids, and the water model have to be accurate. Shortcomings of the force fields, for instance in reproducing the experimental surface tension of water, will lead to deviations in the packing such that simulations can fail to reproduce experimental surface pressure-area isotherms in simple monolayer systems^15–18^. Consequently, consistent methods are required to correctly assign the protonation degree and the packing of lipids to yield robust and meaningful results. In this respect the combination of all-atom MD simulations and X-ray scattering techniques offer promising perspectives, as the latter are uniquely suited to determine the packing of lipid layers^19–21^ and the protonation degree^22^ by quantifying the interfacial counterion excess^23,24^.

The aim of this current work is to introduce a methodology of combining total reflection X-ray scattering and X-ray fluorescence experiments to yield a reliable prediction as benchmark for consistent modeling. We focus on a lipid monolayer composed of the phospholipid POPC and the transfection lipid MC3, one of the most widely used ionizable lipids due to its high transfection efficiency^25^. These simple monolayers, bound to the air/water interface at controlled packing density, are an ideal starting point since they allow us to quantitatively compare the results from all-atom MD simulations and scattering experiments. The joint approach allows us to optimize the protonation degree and the lipid packing. In turn, the simulations yield a detailed insight into the layer structure at the molecular level.

## II. MATERIALS AND METHODS

### A. Chemicals

The transfection lipid DLin-MC3-DMA (MC3, see Fig. 1) was purchased from MedChemExpress (Monmouth Junction, NJ, USA). The phospholipid 1-palmitoyl-2-oleoyl-glycero3-phosphocholine (POPC, see Fig. 1) was purchased from Sigma-Aldrich (Merck KGaA, Germany). Potassium bromide (KBr), Hydrobromic acid (HBr) 48 wt.% in H_2_O and Tris(hydroxymethyl)-aminomethan (Tris-Base, Cl^*−*^-free) were of analytical grade, purchased also from Sigma-Aldrich (Merck KGaA, Germany) and used without further purification. Chloroform (purity*≥* 99%, anhydrous) and methanol (purity*≥* 99.9%) were purchased from Sigma-Aldrich (Merck KGaA, Germany) and used as received.

**FIG. 1.**
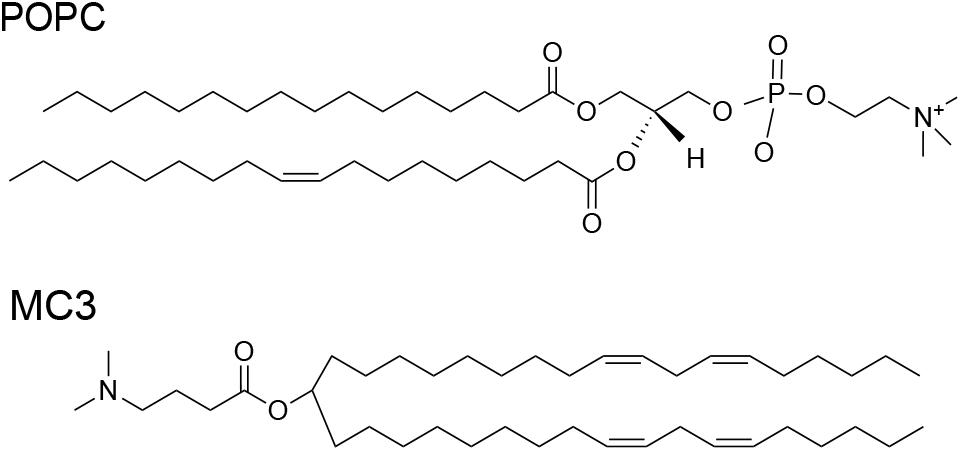
Molecular structures of the phospholipid POPC and the ionizable lipid MC3.

### B. Sample preparation

The mixture of MC3-POPC (20:80 mol%) was prepared by mixing the two dry components and subsequently dissolving them at a concentration of (1 mg/ml) in chloroform-methanol (7:3 v/v). This solution was then spread onto an air-aqueous interface. The resulting monolayer film was laterally compressed until reaching a final lateral pressure of *π* = 30 mN/m while in some cases recording isotherms (see Fig. S1 in the Supporting Information). Two solutions were used as subphases for different experiments: [KBr 2 mM, Tris-Base 5 mM, pH 7.5] and [KBr 2 mM, Tris-Base 5 mM, pH 5.0. The corresponding pHs were reached by adding few drops of a 1M HBr solution. The resulting Br^*−*^ concentration after titration was 5.8 mM for pH 7.5 and 6.8 mM for pH 5.0 (see Supporting Information for further details). We note that Tris-Base does not buffer very well anymore at pH 5.0, but sufficiently well to stabilize the pH over the time of the measurement (*≈* 1 h). We worked at comparatively low Br^*−*^ bulk concentrations in order to minimize undesired X-ray fluorescence contributions from the illuminated aqueous bulk region close to the surface.

### C. Grazing-incidence X-ray scattering techniques

Grazing-incidence X-ray scattering experiments (GIXOS and TRXF, see below) were carried out at the beamline P08 at storage ring PETRA III of Deutsches Elektronen-Synchrotron (DESY, Hamburg, Germany), with the settings described in references^26,27^, from which the following text is partially reproduced. The Langmuir trough (Riegler & Kirstein, Potsdam, Germany) was located in a hermetically sealed container with Kapton windows, and the temperature was kept at 20 ^*°*^C by a thermostat. The container was constantly flushed with a stream of humidified helium (He) to prevent air scattering and the generation of reactive oxygen species. The synchrotron Xray beam was monochromatized to a photon energy of 15 keV, corresponding to a wavelength of *λ* = 0.827 Å. The incident angle was adjusted to *θ*_*i*_ = 0.07 ^*°*^, slightly below the critical angle of total reflection, *θ*_*c*_ = 0.082 ^*°*^. A ground glass plate was placed less than 1 mm beneath the illuminated area of the monolayer in order to reduce mechanically excited surface waves. The beam footprint on the water surface was 1 mm x 60 mm as imposed by the incident beam optics.

Under total-reflection conditions, an X-ray standing wave (SW) is formed at the air/water interface. The penetration depth Λ of its evanescent tail into the aqueous hemispace is a function of the angle of incidence *θ*_*i*_^28^ and for the incident angle used is ≃ 8 nm. The exact shape *φ* (*z*) of the SW intensity along the vertical position *z* (for a given incident angle) follows from the interfacial electron density profile *ρ*_e_(*z*), and can be calculated as described previously^21,29^.

#### 1. Grazing-incidence X-ray off-specular scattering (GIXOS)

Analogous to conventional X-ray reflectometry, GIXOS allows reconstructing the interfacial electron density profile (i.e., the laterally-averaged structure of the lipid layer in the direction perpendicular to the surface) from the *q*_*z*_-dependent scattering intensity, however at fixed incident angle. The details of this technique are described in reference^20,30,31^. As explained more briefly in Kanduc et al.^32^, the *q*_*z*_-dependence of the diffuse scattering intensity *I*(*q*_*xy*_ ≠ 0, *q*_*z*_) recorded at lowenough yet finite *q*_*xy*_ (“out of the specular plane”) with the help of a narrow slit contains information equivalent to that of the conventional reflectivity *R*(*q*_*z*_) and can be transformed as:

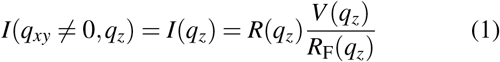

to good approximation, where *R*_F_(*q*_*z*_) the reflectivity of an ideal interface between the two bulk media and *V* (*q*_*z*_) is the Vineyard function^33^. The approximation is based on the assumption of conformal topographic roughness of all surfaces, which is justified for a lipid monolayer subject to capillary wave roughness. In the present work, the GIXOS signal was measured at *q*_*xy*_ = 0.04 Å^*−*1^.

Like conventional reflectivity curves, GIXOS curves can be computed on the basis of an assumed interfacial electron density profile *ρ*_e_(*z*), via the phase-correct summation of all reflected and transmitted partial waves occurring at the density gradients^21,29^. Here, we either modeled *ρ*_e_(*z*) as a set of rough slabs representing the tail and headgroup regions of the monolayer with adjustable thickness, electron density, and roughness parameters^27,31,32^ or extracted *ρ*_e_(*z*) directly from the MD simulations. In all cases, *ρ*_e_(*z*) was then discretized and the corresponding *q*_*z*_-dependent reflectivities *R*(*q*_*z*_) were calculated from the Fresnel reflection laws at each slab-slab interface using the iterative recipe of Parratt^34^ or the equivalent Abebe’s matrix method^35^ and subsequently multiplied with *V* (*q*_*z*_)*/R*_F_(*q*_*z*_) to obtain the theoretical GIXOS signal. For fits to the experimental data, a constant scale factor and a constant intensity background was applied to the modeled intensities, and the fitting range was limited to the consensus range of validity 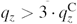, where 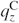 is the *q*_*z*_-value at the critical angle of total reflection)^31^.

#### 2. Total-reflection X-ray fluorescence (TRXF)

The fluorescence signal induced via photoelectric ionization by the X-ray beam under total reflection conditions was recorded with an Amptek X-123SDD detector (Amptek, Bedford, USA). The detector was placed almost parallel to the water surface and perpendicular to the X-ray beam axis, in order to keep elastic and Compton scattering into the detector as low as possible. The center of the detector view angle was set to coincide with the beam footprint position on the water surface.

The fluorescence intensity *I*_F_ emitted by the Br^*−*^ counterions present at the interface is determined by their interfacial depth profile *c*(*z*)^36^. On a quantitative level, *I*_F_ is proportional to the spatial integral over the product of *c*(*z*) and the known SW intensity profile *φ* (*z*) introduced above,

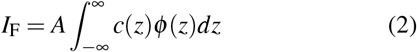

where the prefactor *A* can be calibrated with a suitable reference. Here, we used the surface of a bare KBr solution as reference, for which *c*(*z*) is known and can be considered constant^21,36^. Experimentally, *I*_F_ was obtained by fitting the intensity peak associated with the respective *K*_*α*_ or *L*_*β*_ emission lines of Br (at *≈*12.0 keV and *≈*1.5 keV) in the recorded fluorescence spectra with a Gaussian function. Constraints on the peak positions were imposed based on the tabulated line energies^37^. The standing wave *φ* (*z*) was calculated with a slab-model representation of the interfacial electron density profile^21^, whose parameters were previously obtained in fits to the GIXOS curves. Note that roughness can be neglected for the computation of *φ* (*z*) when *q*_*z*_ is low, as is the case under total reflection^36^.

### D. Theoretical predictions of the protonation degree

#### 1. Infinite dilution limit

The ionizable MC3 lipids are protonated according to the reaction

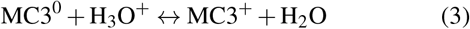

At infinite dilution of the lipids, the protonation degree is given by the Henderson-Hasselbalch equation

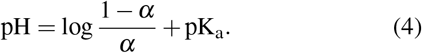

for the protonation degree *α*, where we assume pK_a_ = 9.47 at infinite dilution^4^.

#### 2. Self-consistent equation for the protonation degree

The pK_a_ is related to the equilibrium constant K_a_ by

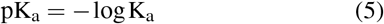

At the monolayer/water interface, the protonation of MC3 lipids are influenced by the distribution of H_3_O^+^ ions and the equilibrium constant K_a_ is given by

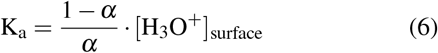

where [H_3_O^+^]_surface_ is the concentration at the position of the ionizable group in the monolayer. The surface concentration is related to the bulk concentration via a Boltzmann distribution^38^

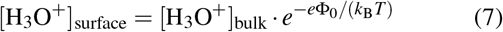

where *e* is the elementary charge, *k*_B_*T* the product of Boltzmann constant and temperature, and Φ_0_ the electrostatic potential at the position of the ionizable group. Combining Eqs. 6-7 and using the definitions for pH and pK_a_ yields a self-consistent equation for the pH as a function of the protonation degree^39^

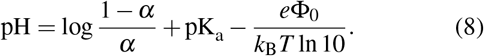

The last term is the electrostatic contribution that takes the H_3_O^+^ distribution dependent on the surface potential into account. Without this term the ordinary equation for the acid deprotonation equilibrium in bulk solution is obtained (Eq. 4). The electrostatic potential Φ(*z*) is calculated from the Poisson Boltzmann equation at a planar interface

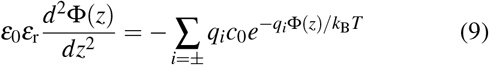

Here, *z* is the distance perpendicular to the surface, *q*_*±*_ = ± *e* is the ion charge in a 1:1 electrolyte, *c*_0_ is the bulk salt concentration, *ε*_0_ is the dielectric constant of vacuum, *ε*_r_ is the relative dielectric constant of water, and Φ(*z*) is the electrostatic potential. For the constant charge boundary condition, we obtain the analytical solution

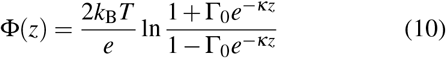

where Γ_0_ is related to the surface potential via

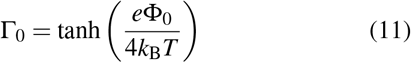

and *κ*^*−*1^ is the Debye screening length

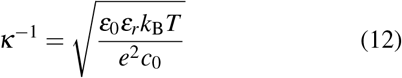

The surface potential Φ_0_ = Φ(0) is obtained from

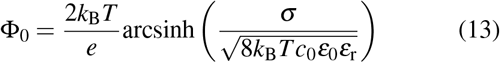

The surface charge density *σ* is related to the area per lipid via *σ* = *αe f*_MC3_*/A*_lip_, where *f*_MC3_ is the fraction of MC3 among all lipids and *e* is the elementary charge. The experimentally obtained ion concentrations of *c*_0_ = 6.8 mM for *α* = 67.5 % and *c*_0_ = 5.8 mM for *α* = 15 % were used for the theoretical predictions. In the self-consistent approach, the protonation degree depends on the surface potential Φ_0_. Clearly, Φ_0_ depends on *σ* and hence on the protonation degree *α* and the area per lipid *A*_lip_. In our approach, Φ_0_ and *σ* were obtained from the results of the final simulations with optimized *α* and *A*_lip_ based on the experiments. *σ* was obtained by dividing the total number of protonated MC3 lipids in one leaflet of the monolayer by the area of the leaflet. The values of *σ* calculated at *α* = 67.5 % and *α* = 15 % were 0.029 Cm^*−*2^ and 0.006 Cm^*−*2^, respectively. With these values, we obtain surface potentials of Φ_0_ = 92.5 mV and Φ_0_ = 33.5 mV at *α* = 67.5 % and *α* = 15 %, respectively.

### E. MD Simulations

We simulated the MC3-POPC monolayers (20:80 mol %) using the Gromacs simulation package (v-2021.5)^40^. The monolayers were created by translating the leaflets of an equilibrated bilayer which in turn was created using the MemGen webserver^41^. The AMBER Lipid 17 force field^42^ was used to describe the POPC lipids. For the cationic and neutral MC3 molecules we used our recently developed force fields^43^. These parameters have the advantage that they closely reproduce the structure of MC3 in lipid layers as judged by neutron reflectometry experiments. In addition, the parameters are compatible with the AMBER force field family. Ions were described using the Mamatkulov-Schwierz force field parameters^44^ and the TIP3P water model was used^45^. The systems contained 160 POPC and 40 MC3 lipids per monolayer, and 60 water molecules per lipid. Chloride ions were added to neutralize the positive charge of the protonated MC3 lipid molecules. The systems were energy minimized using a gradient descent algorithm. The temperature of the systems was maintained at 293.15 K using the velocity rescaling thermostat with stochastic term^46^ and a time constant of 1.0 ps. For the *NPT* simulations, semi-isotropic Parrinello-Rahman^47^ was used. The Van der Waals interaction potential was cut off and shifted to zero at 1.2 nm. Electrostatic interactions were cut-off at 1.2 nm and the particle mesh ewald (PME) method^48^ with a fourier grid spacing of 0.12 nm was used to evaluate the long-range electrostatics part. All bonds involving hydrogens were constrained using the LINCS algorithm^49^. A timestep of 2.0 fs was used to integrate the equations of motion. Analysis was performed using Gromacs inbuilt routines and with MDAnalysis^50^ and trajectories were visualized with VMD^51^. To obtain the desired area per lipid, the in-plane components *P*_*xx*_ and *P*_*yy*_ of the pressure tensor were varied in 11 steps from -50 bar to -34 bar. These simulations had durations of 100 ns. A semi-isotropic barostat was employed, such that *P*_*xx*_ = *P*_*yy*_. The box size in *z*-direction, *L*_*z*_ = 20 nm, was kept constant. The surface tension *γ* can be deduced from the pressure tensor components^52^ as

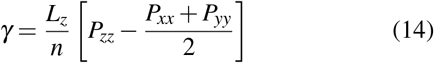

where *n* = 2 is the number of interfaces in the simulation box^53^. The value of *γ* was directly obtained using the Gromacs gmx energy routine. The covered range of in-plane pressure tensor components corresponds to a surface tension range of 34 mN/m ≤*γ* ≤50 mN/m. The surface pressure is calculated from *π* = *γ*_0_− *γ*, where *γ*_0_ = 58 mN/m is the surface tension of TIP3P water at 293.15 K as obtained through extrapolation of the data by Vega et al.^52^. The last 50 ns of each of these simulation at given value of surface pressure was used to obtain the electron density, from which the GIXOS curves were evaluated using Eq. 1. The Refnx python package^54^ was used to obtain the reflectivity profiles (*R*(*q*_*z*_) and *R*_F_(*q*_*z*_) in Eq. 1). The simulations which best reproduced the experimental GIXOS curves at pH 5.0 and pH 7.5 for *α* = 1.0 had the parameters summarized in Table I. The simulations with experimentally validated values of *A*_lip_ and *α* were performed for 500 ns in the *NVT* ensemble and had the parameters summarized in Table II.

**TABLE I.** Simulation parameters that best reproduce the experimental GIXOS curves for *α* = 1.0.

| pH | Fixed Parameters |  |  | Calculated Parameters |  |  |  |  |
| --- | --- | --- | --- | --- | --- | --- | --- | --- |
| | $\alpha$ | $L_z$<br>[nm] | $P_{xx}/P_{yy}$<br>[bar] | $P_{zz}$<br>[bar] | $L_x/L_y$<br>[nm] | $A_{lip}$<br>[nm<sup>2</sup>] | $\gamma$<br>[mN/m] | $\pi$<br>[mN/m] |
| 5.0 | 1.0 | 20.0 | 46.0 | $-0.36 \pm 0.72$ | $12.10 \pm 0.02$ | $0.73 \pm 0.002$ | $45.9 \pm 0.9$ | $12.1 \pm 0.9$ |
| 7.5 | 1.0 | 20.0 | 48.0 | $1.26 \pm 1.0$ | $12.25 \pm 0.02$ | $0.75 \pm 0.003$ | $49.8 \pm 1.2$ | $8.2 \pm 1.2$ |

**TABLE II.** Parameters used for the simulations with experimentally validated area per lipid and protonation degree.

| pH | Fixed Parameters |  |  |  | Calculated Parameters |  |  |  |  |
| --- | --- | --- | --- | --- | --- | --- | --- | --- | --- |
| | $\alpha$ | $L_x/L_y$<br>[nm] | $L_z$<br>[nm] | $A_{lip}$<br>[nm<sup>2</sup>] | $P_{xx}$<br>[bar] | $P_{yy}$<br>[bar] | $P_{zz}$<br>[bar] | $\gamma$<br>[mN/m] | $\pi$<br>[mN/m] |
| 5.0 | 0.675 | 12.10 | 20.0 | 0.73 | $-47.8 \pm 0.35$ | $-48.2 \pm 0.4$ | $-0.36 \pm 0.46$ | $47.6 \pm 0.6$ | $10.4 \pm 0.6$ |
| 7.5 | 0.150 | 12.25 | 20.0 | 0.75 | $-48.8 \pm 0.6$ | $-49.2 \pm 0.8$ | $-0.42 \pm 0.56$ | $49.1 \pm 0.8$ | $8.9 \pm 0.8$ |

## III. RESULTS AND DISCUSSION

The lipid monolayers contain 80 mol% of the charge-neutral zwitterionic phospholipid POPC and 20 mol% of the ionizable lipid MC3 (Fig. 1). Dependent on the pH, the ionizable MC3 can be neutral (MC3^0^) or positively charged (MC3^+^) such that the monolayer contains up to three lipid species.

Our aim is to elucidate the influence of pH on the properties of the monolayers at two different pH values, pH 7.5 and pH 5.0, representing the extracellular and the endosomal milieu, respectively. However, to perform simulations two important parameters have to be chosen. The first one is the lateral monolayer packing in terms of the average area per lipid, *A*_lip_. It should match the experimental one at a lateral pressure of *π* = 30 mN/m, a value representative of the packing in lipid bilayers^55^. The second one is the protonation degree *α* of MC3, which should be consistent with the experimental value at given pH conditions. To correctly assign *A*_lip_ and *α*, we obtain complementary information from two synchrotron-based grazing-incidence X-ray scattering techniques, namely grazing-incidence X-ray off-specular scattering (GIXOS) and total-reflection X-ray fluorescence (TRXF)^36,56,57^. While GIXOS, like conventional reflectometry, precisely reveals the monolayer thickness^27^ (albeit with limited insights into the conformations and lateral distributions of the different molecular species), TRXF is highly sensitive to the interfacial accumulation of counterions at charged monolayers^23,24,36^ and can therefore serve to determine protonation degrees^22^. As will be shown later, the accuracy in the experimental determination of *α* is further enhanced by including the counterion distribution from the simulations.

Fig. 2 outlines the methodology to obtain consistent values for *A*_lip_ and *α*. Initially, *α* is estimated based on the Henderson-Hasselbach equation which predicts *α* = 1 for both pH values (Fig. 3). Note that the self-consistent equation yields an improved estimate but requires the surface charge density, which is not known a-priori. In a second step, we performed simulations with the assumed *α* value while systematically varying *A*_lip_ to identify the values that match the layer thickness obtained in the GIXOS measurements at the two pH conditions. In the next step, the pH-dependent protonation degrees *α* are deduced from the TRXF measurements while imposing the known values of *A*_lip_ as well as the shape of the counterion distribution found in the simulations. Finally, new simulations with the experimentally validated values of *A*_lip_ and *α* are performed for both pH conditions. In the following, we discuss the results from each step in detail.

**FIG. 2.**
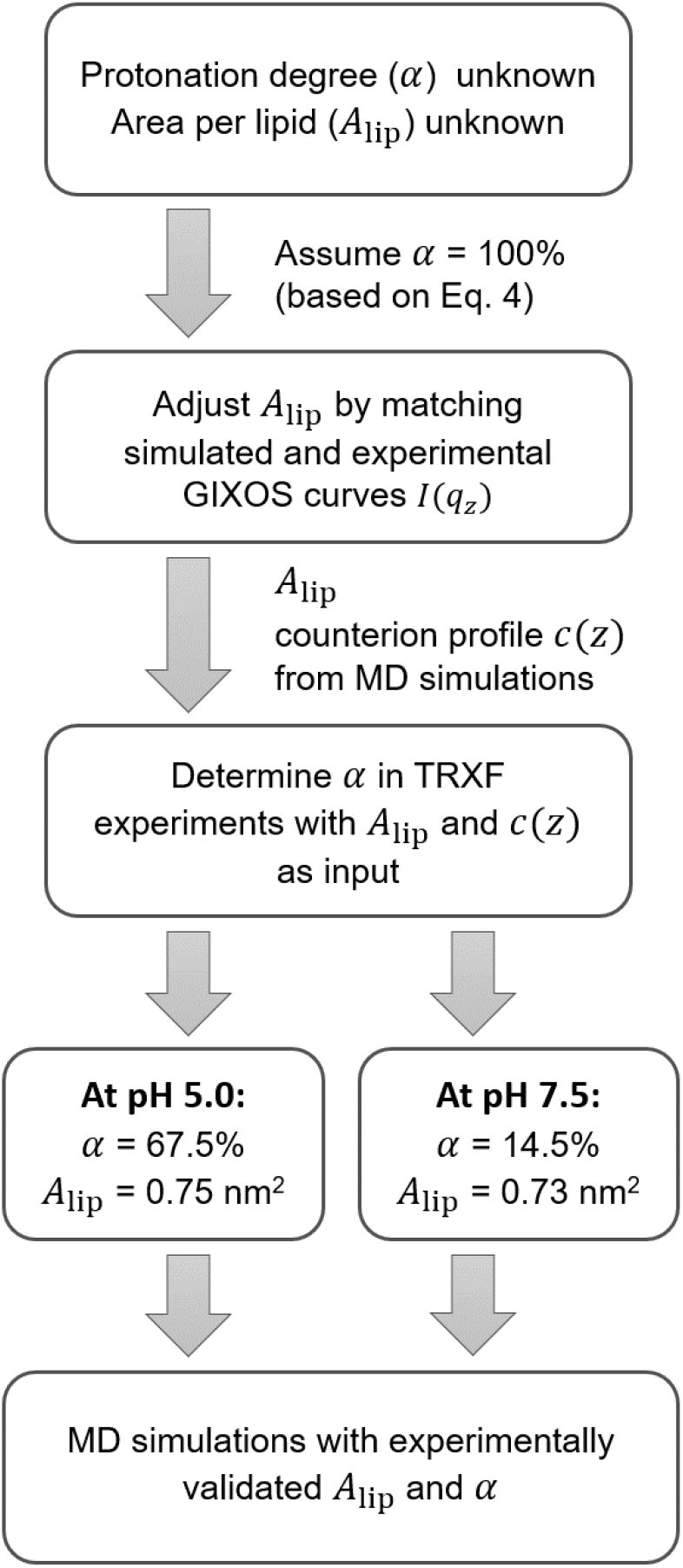
Methodology of combining all-atom molecular dynamics simulations with experimental total reflection X-ray scattering and X-ray fluorescence for consistent modeling of the area per lipid and the protonation degree.

**FIG. 3.**
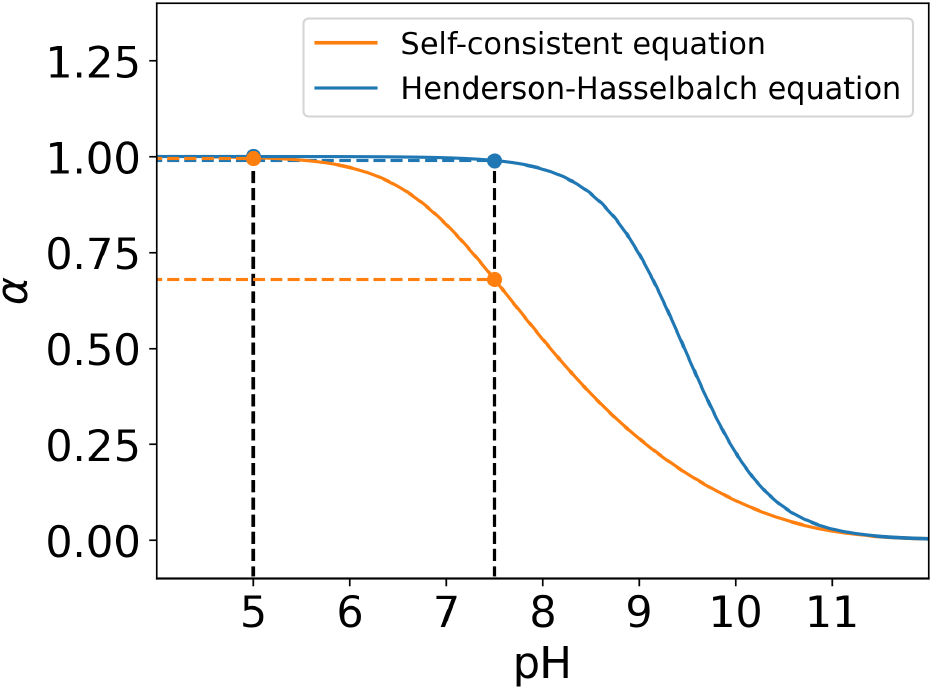
Dependence of the protonation degree *α* on pH obtained from the Henderson-Hasselbach equation 4 and the self-consistent equation 8 using pK_a_ = 9.4^4^. Dashed straight lines indicate the predictions for the two experimental pH values, 5.0 and 7.5.

### A. Determination of the area per lipid

Fig. 4 A shows the GIXOS curves obtained experimentally with monolayers at *π* = 30 mN/m on aqueous subphases with pH 7.5 (top) and with pH 5.0 (bottom). The subphases additionally contained defined concentrations of Br^*−*^ and K^+^ ions, as required for the simultaneous TRXF measurements discussed further below. The curves contain information on the electron density profiles of the monolayers, where the overall layer thicknesses are mainly encoded in the *q*_*z*_-positions of the intensity minima. The lines superimposed to the experimental data points are the simulated GIXOS curves computed from the electron density profiles obtained in simulations. Based on the Henderson-Hasselbalch equation (Eq. 4) the protonation degree is *α≈* 1 (Fig. 3). However, significant deviations from the experimental curve (dashed line in Fig. 4 A) are obtained with simulations at the experimental surface pressure corresponding to a surface tension of *γ* = *γ*_0_− *π* = 28 mN/m, with *γ*_0_ = 58 mN/m for TIP3P water (see Methods section). Clearly, the packing of the lipids and hence *A*_lip_ is not reproduced. This deviation is likely caused by the TIP3P water model which, like most water models, does not yield the correct surface tension of water^18,52^. In order to provide improvement, we systematically vary *A*_lip_ in the simulations to obtain the best-matching value for each pH, characterized by the minimum *χ*^2^ deviation (Figs. S4 and S5) from the experimental GIXOS data. The simulated GIXOS curves obtained after optimization (solid lines) reproduce the experimental data well, demonstrating that the simulations properly capture the overall structure of the monolayers.

**FIG. 4.**
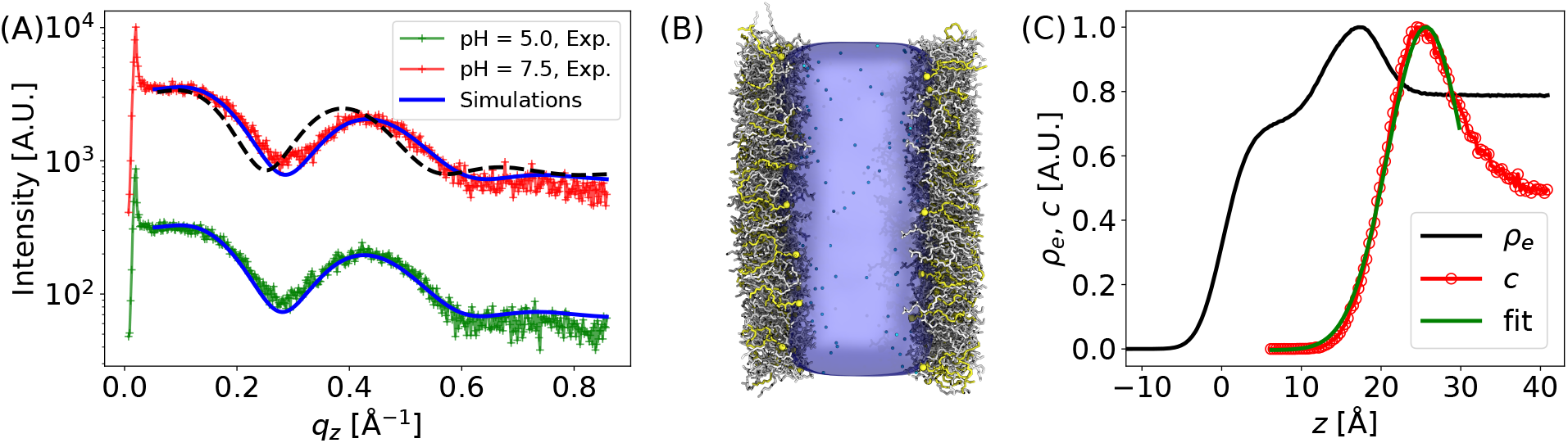
(A) Comparison of experimental GIXOS curves with the simulated ones for the optimized values of *A*_lip_. The system consists of 20 % MC3^+^ and 80 % POPC. The curves at different pH values are vertically shifted for clarity. The dashed black line is obtained by using the same surface pressure as in the experiments (*π* = 30 mN/m) and corresponds to *A*_lip_ = 0.63 nm^2^. The solid blue lines correspond to the simulations with optimized area per lipid (*A*_lip_ = 0.75 nm^2^ for pH = 7.5 and *A*_lip_ = 0.73 nm^2^ for pH = 5.0. (B) Snapshot from a simulation with *A*_lip_ = 0.63 nm^2^ and *α* = 1 after 100 ns of equilibration. All other run parameters are summarized in Table I. MC3^+^ is represented in yellow with the head group nitrogen highlighted by yellow spheres. POPC is shown in white and chloride ions in cyan color. (C) Corresponding electron density profile *ρ*_e_(*z*). The plot also shows the counterion density profile *c*(*z*) obtained in the simulations (data points). The solid line superimposed is an analytical fit with Eq. 15.

For pH 7.5 and pH 5.0, the best-matching values are *A*_lip_ = 0.75 nm^2^ and *A*_lip_ = 0.73 nm^2^, respectively, which is in the typical range for lipid bilayers the fluid phase^58^. The associated surface pressures in the simulations are as low as *π ≈* 12 mN/m (pH 5.0, see Table I) and *π ≈* 8 mN/m (pH 7.5, see Table I) and deviate by a factor of *≈* 3 from the experimental value. Interestingly, the agreement is much better on the level of the surface tension, where the simulations yield *γ ≈* 46 mN/m*≈* (pH 5.0, see Table I) and *γ* ≈ 50 mN/m*≈* (pH 7.5, see Table I), compared to the experimental value of *γ* = *γ*_0_*− π* = 42 mN/m, where *γ*_0_ = 72 mN/m. As seen in Tables I and II, the protonation degree of MC3 only has a minor influence on *γ* and *π* for a fixed area per lipid. In conclusion, the lateral density in terms of *A*_lip_ appears to be the most robust parameter for the experimental validation of MD simulations of lipid monolayers.

The electron density profile *ρ*_e_(*z*) and the counterion density profile *c*(*z*) obtained in the simulations for *A*_lip_ = 0.73 nm^2^ are shown in Fig. 4 C. The profiles are consistent with earlier reports on fluid lipid monolayers at comparable lateral pressures^19^, however the electron density maximum representing the lipid headgroups is slightly less pronounced than for commonly studied phospholipid layers, because MC3 has a more compact headgroup without the electron-rich P atom.

### B. Determination of the protonation degree

The pH-dependent protonation degree of MC3 was determined experimentally through measurements of the interfacial excess of Br^*−*^ counterions, Γ, which coincides with the density of positive net charges in the monolayer^36^. Br^*−*^ rather than the more common Cl^*−*^ was used as counterion in these experiments because the used TRXF setup was slightly more sensitive to heavier elements. Note, however, that Br^*−*^ is equally suited as indicator of the protonation degree. Fig. 5 A shows the X-ray fluorescence spectra in a narrow energy range around the peak associated with the *K*_*α*_ emission line of Br. The spectra were obtained under total reflection conditions with the bare aqueous subphases and with lipid monolayers at *π* = 30 mN/m both at pH 5.0 and pH 7.5. The complete spectra are shown in the Supporting Information (Fig. S2). The intensities of the emission lines are much higher in the presence of the monolayer at both pH values, which reflects the accumulation of Br^*−*^ ions at the interface required to compensate the positive charges of MC3^+^. The *L*_*β*_ emission lines behave consistently (see Supporting Information). The intensity is much higher at pH 5.0 than at pH 7.5, evidencing that the positive charge density is much higher at the lower pH and, thus, the protonation degree *α*. This observation immediately implies that *α* at pH 7.5 is much lower than 100 %, in contradiction to the estimation from the Henderson-Hasselbalch equation. The slight difference in the intensities observed with the bare subphases at the two pH values can be attributed to the slightly different concentration of Br^*−*^ as a result of pH titration with HBr (see Methods section).

**FIG. 5.**
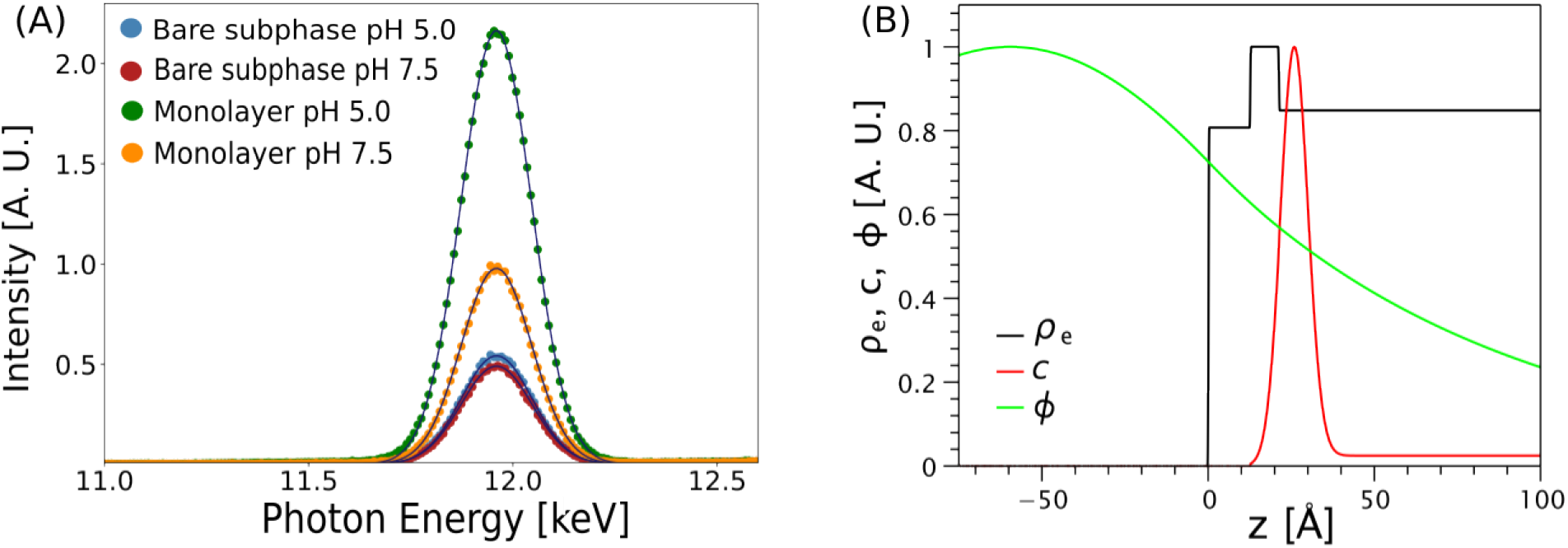
(A) X-ray fluorescence spectra around the *K*_*α*_ emission peak of the Br^*−*^ counterions obtained with the bare aqueous subphases and with lipid monolayers at two pH values. (B) Illustration of the model used for data analysis (here pH 5.0): the two-slab representation of the experimentally obtained electron density *ρ*_e_(*z*), the standing wave intensity *φ* (*z*), and the counterion profile *c*(*z*). For the sake of clarity, all curves are normalized to have maximal values of 1.

The following quantitative analysis takes the counterion profiles *c*(*z*) from the MD simulations into account. This approach can be viewed as a refinement with respect to commonly used treatments of the counterion distribution in terms of thin adsorption layers^27,36^ or related approximations^59^. Here, the simulation-based distributions were modeled as a (truncated) Gaussian function on top of the constant experimental bulk concentration *c*_0_,

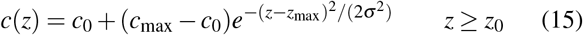

where *z*_0_ defines the interface between the tail and headgroup slabs. This functional form was fitted to the numerical counterion profiles extracted from the simulations (see solid line in Fig. 4 C), yielding *z*_max_ = *z*_0_ + 0.56 nm; *σ* = 0.48 nm for pH 5.0 and *z*_max_ = *z*_0_ + 0.53 nm; *σ* = 0.47 nm for pH 7.5. With that, *c*_max_ remains as the only adjustable parameter, which is varied so as to match the measured fluorescence intensities *I*_F_ in the presence of the lipid monolayer and at the given pH, according to Eq. 2. For illustration, Fig. 5 B displays all ingredients required for this calculation: the two-slab model of the electron density *ρ*_e_(*z*), obtained from the experimental GIXOS curves as described in the Supporting Information (Fig. S3), the corresponding standing wave intensity *φ* (*z*), and the parameterized counterion profile *c*(*z*). The counterion excess then simply follows as

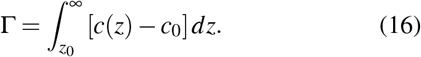

We obtain Γ = 0.185 nm^*−*2^ for pH 5.0 and Γ = 0.039 nm^*−*2^ for pH 7.5. For a known counterion excess, the protonation degree *α* of MC3 follows as:

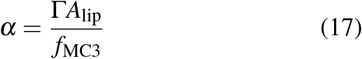

where *f*_MC3_ = 0.2 is the MC3 mole fraction. With that, we obtain *α* = 67.5 % for pH 5.0 and *α* = 14.5 % for pH 7.5. The obtained protonation degrees are significantly lower compared to the prediction from the Henderson-Hasselbalch equation. This is not surprising since the equation is valid at infinite dilution. In the monolayer system, interactions between polar lipids, charged lipids, ions and protons contribute to the protonation degree. The effect of the proton distribution is taken into account in the self-consistent equation 8. The protonation degrees predicted in this way are *α* = 99 % for pH 5.0 and *α* = 68 % for pH 7.5. Although these values provide a more meaningful estimate, they are still significantly higher than the ones experimentally observed and require an a-priori estimate of the surface charge density. To provide improvement, the interactions between the lipids in the monolayer as well as conformational rearrangement and partial dehydration have to be taken into account. In particular the effect of partial dehydration is expected to have a significant effect since the uncharged headgroups are buried while the charged headgroups are solvent exposed (Fig. 6 A and B). In any case, reliable experimental measurements of the protonation degree are imperative for validation and improvement of theoretical predictions.

**FIG. 6.**
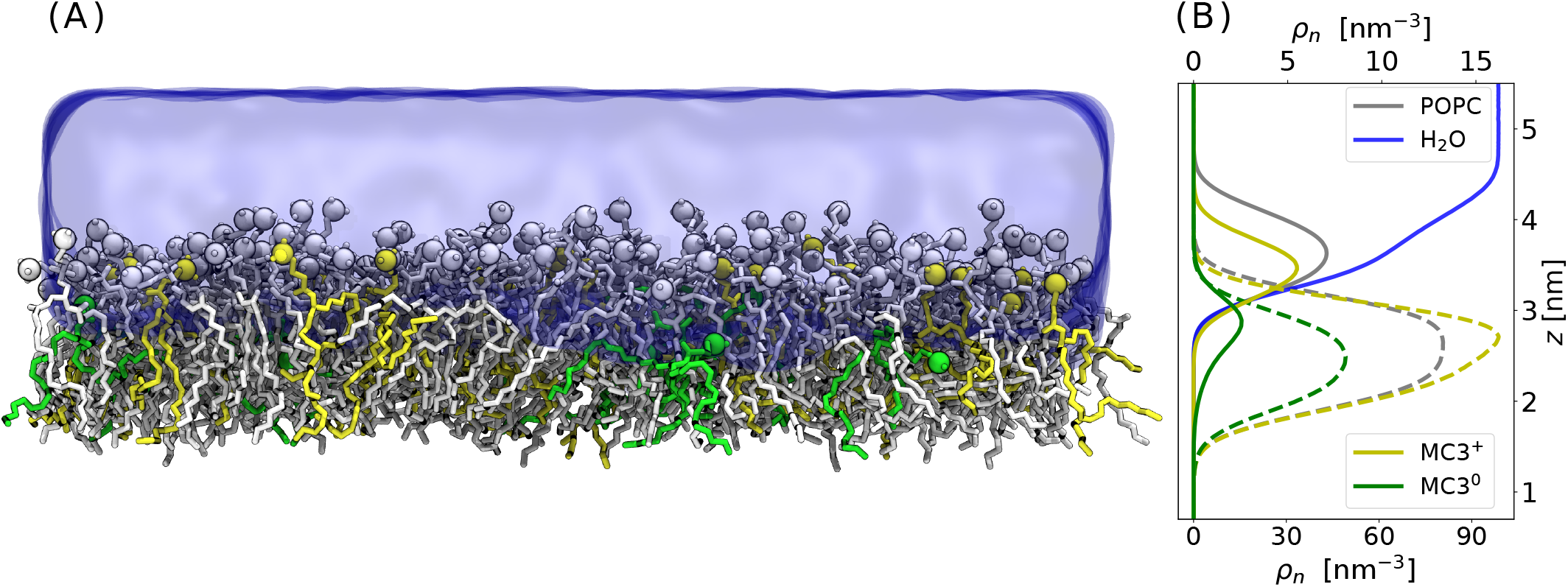
Simulations with experimentally validated *A*_lip_ and *α*. (A) Side view snapshot of one of the monolayers with *α* = 67.5 %. Colors: green (MC3^0^), yellow (MC3^+^) and gray (POPC). (B) Number density profiles *ρ*_n_(*z*) of water and of the head and tail groups of MC3^0^, MC3^+^, and POPC. Solid lines represent head groups and dashed lines represent tail groups. Dotted line: water. The top scale applies to MC3, the bottom scale to POPC and water.

Finally, it is important to confirm that the simulations with the experimentally validated protonation degree still reproduce the experimental GIXOS curves and counterion profiles that were used in the validation procedure (Fig. 2). Otherwise additional iterations would be required. The results show that after only one iteration the results converged and the simulated GIXOS curves do not significantly respond to the updated *α* values (see Fig. S6 in the Supporting Information), and the peak in the counterion distribution still has the same position and width (see Fig. S7 in the Supporting Information).

### C. Simulations with experimentally validated area per lipid and protonation degree

As the last step, MD simulations with the experimentally validated values of *A*_lip_ and *α* for the two pH values were performed, with the parameters summarized in Table II. The correct protonation degrees were imposed by representing the MC3 molecules with suitable numbers of MC3^0^ and MC3^+^ variants. In practice, 6 and 27 out of the 40 MC3 molecules in each monolayer were represented with MC3^+^ for pH 7.5 and pH 5.0, respectively, yielding *α* = 15.0 % and *α* = 67.5 %. These protonation degrees come as close as possible to the experimental values when considering that the numbers of lipids can only be varied in integer steps.

Fig. 6 A shows a snapshot from an equilibrated simulation with *α* = 67.5 %, corresponding to pH 5.0. POPC matrix lipids are represented in white, MC3^0^ in yellow, and MC3^+^ in green. Panel B shows the normalized number density profiles *ρ*_n_ of the tails and headgroups of all lipid species and of water in the *z*-direction, i.e., perpendicular to the membrane plane. The distribution of the positively charged headgroups of MC3^+^ largely overlaps with that of POPC’s zwitterionic headgroups, while the distribution of the uncharged and only weakly polar headgroups of MC3^0^ is strongly shifted towards the hydrophobic region accommodating the chains. As shown in the Supporting information (Fig. S8), consistent observations can be made at *α* = 15.0 %, corresponding to pH 7.5. This behavior has been noticed earlier^43^ and may provide the explanation for the small yet significant increase in *A*_lip_ when increasing the pH from 5.0 to 7.5 (see above). Namely, in contrast to the naive expectation that a reduction in the density of charged lipids would decrease lateral repulsion and thus the lipids’ effective area requirement, the deep insertion of uncharged MC3^0^ into the monolayer effectively increases the area requirement at high pH, where MC3^0^ is more abundant.

We now turn to the in-plane (*xy*) distribution of the three lipid species. Fig. 7 A shows a top-view simulation snapshot of a monolayer after equilibration. Qualitatively, it is seen that both MC3^0^ and MC3^+^, depicted in green and yellow, respectively, distribute rather evenly over the membrane plane, which maximizes their average distance. This behavior is quantified with the help of the normalized in-plane radial distribution functions (RDFs) of the headgroup N atoms, which are shown in panel B. Indeed, while the RDF between two POPC headgroups on approach oscillates around unity and even exhibits a pronounced next-neighbor peak at *r≈* 0.9 nm due to steric and dipole interactions^60^, the RDF between two MC3^+^ headgroups systematically drops below unity on approach, albeit with a few weak oscillations. This drop in the RDF can be attributed to the electrostatic repulsion between the positively charged headgroups. The RDF between two MC3^0^ drops below unity on approach, too, but like-charge repulsion obviously cannot be the reason because MC3^0^ carry no net charge. Calculation of the mean force reveals an oscillatory behavior (Fig. S10). Repulsion occurs at small distances since MC3^0^ leads to a local arrangement of the POPC molecules which oppose to some extent the penetration of other MC3^0^ molecules nearby.

**FIG. 7.**
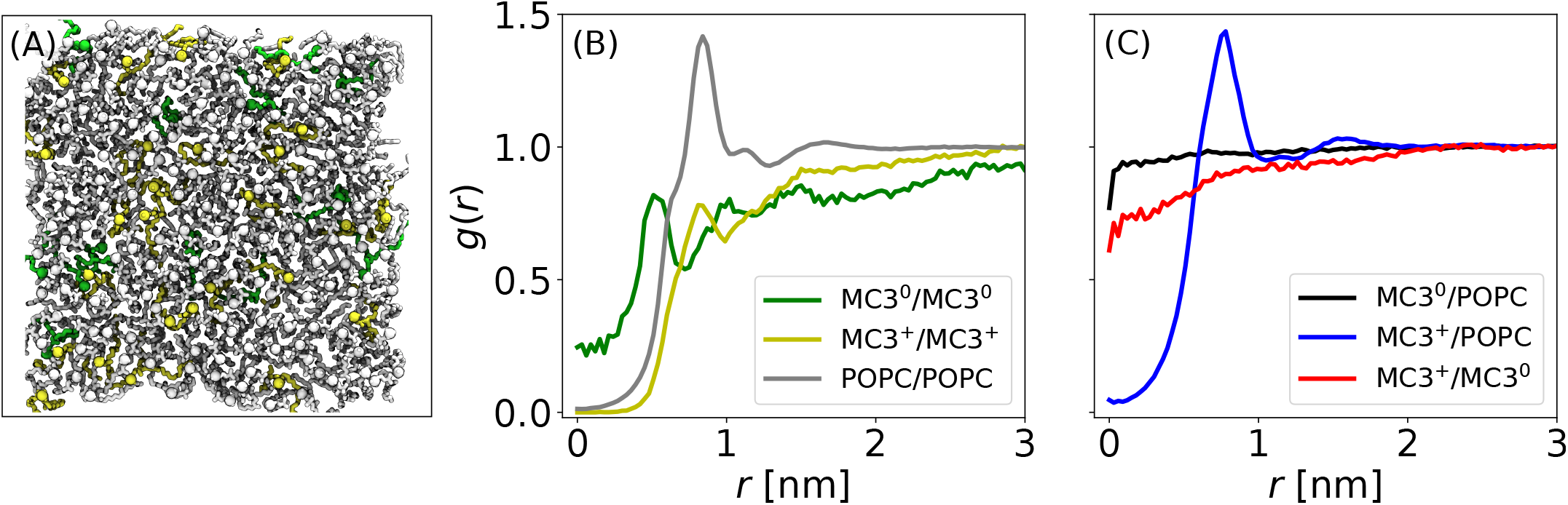
Simulations with experimentally validated *A*_lip_ and *α*. (A) Top view snapshot of one of the monolayers with *α* = 67.5 %. Colors: green (MC3^0^), yellow (MC3^+^) and gray (POPC). (B) In-plane radial distribution functions (RDFs) between the headgroup nitrogens of the same lipid species. (C) RDFs between headgroup nitrogens of different lipid species.

Another qualitative difference between the RDF of two MC3^0^ and the other two RDFs shown in Fig. 7 B is its finite value at *r* = 0. Apparently, two MC3^0^ headgroups can share the same in-plane (*xy*) coordinates because they are somewhat mobile in the third dimension (*z*), even though their depth profile in Fig. 6 B is not significantly broader than those of the other headgroups. To resolve this apparent discrepancy we have calculated the 3D-RDFs of the MC3^0^ headgroups and compared them to the 3D-RDFs of the other headgroups (Fig. S9 in the Supporting Information). It is seen that the radius of mutual steric exclusion is much smaller for MC3^0^ than for the other headgroups, which explains why the probability for two MC3^0^ headgroups to be at the same *xy* position is higher than that for the other headgroups.

Next we take a look at the mixed RDFs shown in Fig. 7 C. Strikingly, the mixed RDFs of MC3^0^ with POPC and of MC3^0^ with MC3^+^ both merely exhibit a weak dip near *r* = 0, demonstrating that there is little steric exclusion because the headgroups of MC3^0^ roam at a different “altitude” than those of POPC and MC3^+^. In contrast, as seen from the mixed RDF of POPC with MC3^+^, their headgroups mutually exclude each other sterically for small *r* and exhibit a pronounced next-neighbor peak at *r≈* 0.9 nm due to attractive interactions between the positively charged MC3^+^ headgroups and the negatively charged phosphate groups in the PC headgroups^61^.

## IV. CONCLUSIONS

Molecular dynamics simulations of ionizable lipids provide detailed insight into their structure but require an a-priori knowledge of the lipid packing and protonation degree. We have introduced a methodology of combining all-atom molecular dynamics simulations with experimental total reflection X-ray scattering and X-ray fluorescence to make a robust and reliable predictions of MC3 lipid area and degree of protonation at pH conditions that mimic the extracellular and endosomal milieus. The combined approach yields a protonation degree *α* = 14.5% and an average area per molecule *A*_lip_ = 0.73 nm^2^ at pH 7.5. At pH 5.0, it yields *α* = 67.5% and *A*_lip_ = 0.75 nm^2^. In both cases the value for the protonation degree differs significantly from the value expected from the Henderson-Hasselbalch equation. Hence, the presetting of the protonation degree in the simulations based on simple theories can fail due to the interactions between polar lipids, charged lipids, ions and protons in complex lipid interfaces.

The simulations with optimized *A*_lip_ and *α*, obtained from the matching methodology, reveal that the headgroups of the uncharged MC3 enter deeply into the hydrophobic monolayer region while the charged MC3 headgroups are solvent-exposed, as seen from the density profiles in Fig. 6. Lipid dehydration upon deprotonation is therefore expected to have a significant effect on the protonation degree and is likely the reason why theoretical predictions based on the HendersonHasselbach equation overestimate the protonation degree for these simple monolayer systems. In addition, the buried uncharged MC3 lipids are laterally almost uncorrelated from the more hydrophilic POPC headgroups and the charged MC3 lipids. This behavior is associated with a non-typical structural response of *A*_lip_ to pH variations, which may play a significant role for the endosomal escape and subsequent biological action of mRNA.

The joint experimental and simulation approach presented here provides a benchmark for consistent modeling. The results can be used to test theoretical models or advanced simulation methods such as free energy simulations or constantpH simulations. The methodology is also suited to investigate changes in protonation and lipid packing in response to interactions with nucleic acids. Another challenge is to extend this methodology for predictions in three-dimensional lipid mesophases and lipid nanoparticles.

## ACKNOWLEDGMENTS

We acknowledge DESY (Hamburg, Germany), a member of the Helmholtz Association HGF, for the provision of experimental facilities. Parts of this research were carried out at PETRA III and we would like to thank Chen Shen for assistance in using P08. Beamtime was allocated for proposal I-11009842. We thank Gerald Brezesinski and Tetiana Mukhina for help with the synchrotron experiments. This research was supported by the Bundesministerium für Bildung und Forschung (BMBF) within the Röntgen-Ångström cluster project Medisoft, number 05K18EZA and 05K18WMA, and by the Emmy Noether program of the Deutsche Forschungsgemeinschaft (DFG, German Research Foundation), number 315221747. We acknowledge the GOETHE HLR for super-computing access. The authors gratefully acknowledge the scientific support and HPC resources provided by the Erlangen National High Performance Computing Center (NHR@FAU) of the Friedrich-Alexander-Universität Erlangen-Nürnberg (FAU) under the NHR project b119ee. NHR funding is provided by federal and Bavarian state authorities. NHR@FAU hardware is partially funded by the German Research Foundation (DFG), number 440719683.

## DATA AVAILABILITY STATEMENT

Experimental and simulation raw data and derived data supporting the findings of this study are available from the corresponding authors upon reasonable request.

