## supplementary text and figures for "Combining molecular dynamics simulations and X-ray scattering techniques for the accurate treatment of protonation degree and packing of ionizable lipids in monolayers"

#### Experiments

##### pH Titration

In order to verify the final concentration of $Br^-$, a solution with 5 mM Tris and 2 mM KBr. The initial MilliQ water volume was 0.5 L. To reach pH 7.5, 1.85 mL of 1 M HBr had to be added, while to reach pH 5.0, 2.3 mL of 1 M HBr had to be added. These quantities were found to be reproducible within an uncertainty of ±0.1 mL. With that, the final concentration of $Br^-$ in the solution was 5.8 mM for pH 7.5 and 6.8 mM for pH 5.0, considering the sum of the titrated concentration and the original 2 mM $[Br^-]$. These numbers were consistent within ±0.2 mM with the theoretical prediction, based on Henderson-Hasselbalch equation.[1,2]

#### Isotherms

Pressure-area isotherms (see Fig. S1) were collected at room temperature with a KSV NIMA Langmuir trough (KSV, Finland). A Wilhelmy paper plate sensor was used as pressure sensor. The lipid solution was spread over the water subphase using a Hamilton syringe and the solvent was let evaporate completely for ≈ 15 min. The monolayers were laterally compressed to the target surface pressure with a compression speed of ≤ 1%/min.

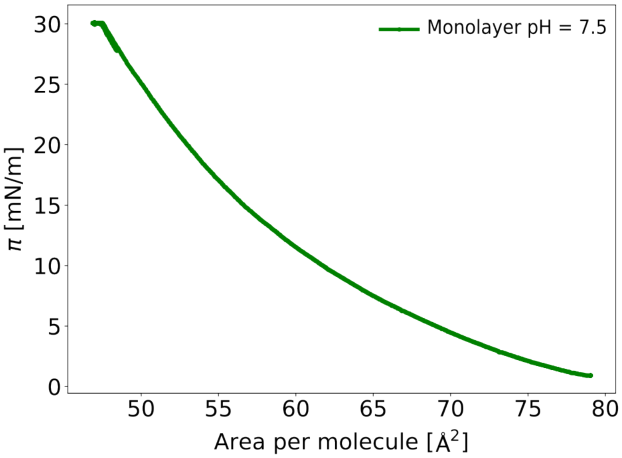


Figure S1: Langmuir isotherm of MC3-POPC (20:80 mol%) mixture at pH 7.5.

#### TRXF spectra

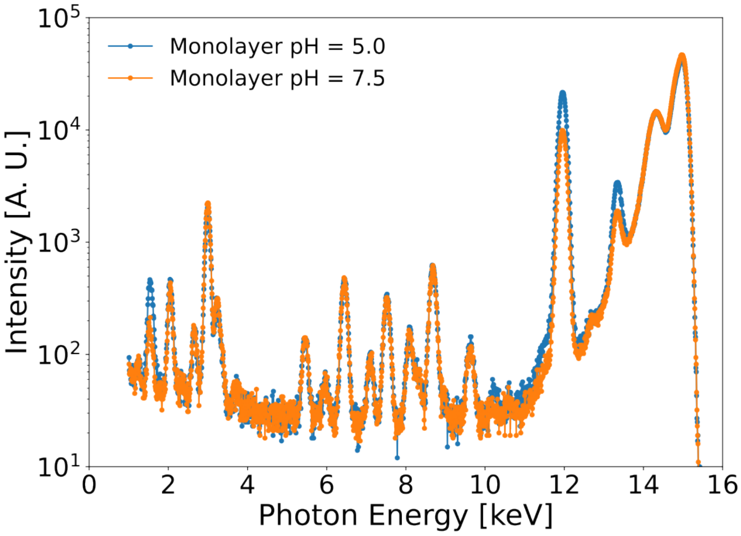


Figure S2: Fluorescence signal induced via photoelectric ionization of MC3-POPC (20:80 mol%) lipid monolayer at $\pi = 30$ mN/m both at pH 5.0 and pH 7.5, recorded with an Amptek X-123SDD detector (Amptek, Bedford, USA).

#### Fitting GIXOS curves using two-slab model

The data were analyzed describing the monolayer film with two homogeneous slabs or boxes of adjustable thickness $d$ and electron density $\rho$, which physically represent different portions of the lipid monolayers: headgroups and hydrocarbon tails. The interfaces between slabs are subject to interfacial roughness to an adjustable extent encoded in the roughness parameters $\sigma$. The experimental data points were fitted with a theoretically modeled GIXOS signal (Fig. S3 A) based on a two-layer description after optimization of the parameters $d$ and $\rho$ of the slabs and of the $\sigma$ parameters for the interfaces. The resulting electron density profiles are shown in Fig. S3 B. The best-matching parameters are summarized in table S1.

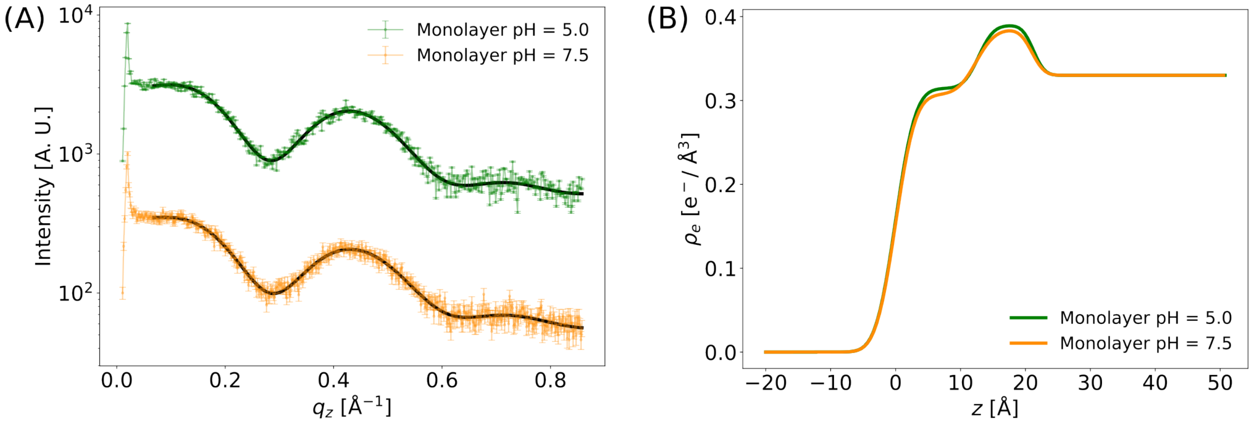


Figure S3: (A) GIXOS data (data points) with fits (solid lines). (B) Corresponding electron density profiles.

Table S1: Slab model parameters obtained for the monolayers at $\pi = 30$ mN/m at pH 5.0 and pH 7.5.

| parameter | pH 5.0 | pH 7.5 |
|---|---|---|
| $\rho_{\text{tail}}$ [$e^-$/Å$^3$] | 0.31 | 0.31 |
| $\rho_{\text{head}}$ [$e^-$/Å$^3$] | 0.39 | 0.38 |
| $\sigma_{\text{air/tails}}$ [Å] | 2.39 | 2.39 |
| $\sigma_{\text{tails/headgroups}}$ [Å] | 1.83 | 2.29 |
| $d_{\text{tail}}$ [Å] | 12.76 | 12.38 |
| $d_{\text{head}}$ [Å] | 8.46 | 8.61 |

### Simulations

#### GIXOS curves from simulations at different values of $A_{\text{lip}}$

To obtain the optimum area per lipid from simulations, the value of $A_{\text{lip}}$ was systematically varied and the corresponding simulated GIXOS curves $I_{\text{sim}}(q_z)$, computed from the associated electron density profiles shown in Fig. S4 A were compared to the experimental GIXOS curves $I_{\text{exp}}(q_z)$, as shown in S4 B. The optimum area corresponds to the minimum in the $\chi^2$ deviation

$$\chi^2 = \sum_{q_z} \frac{(I_{\text{sim}}(q_z) - I_{\text{exp}}(q_z))^2}{N} \tag{1}$$

, where $N$ is the number of data points.

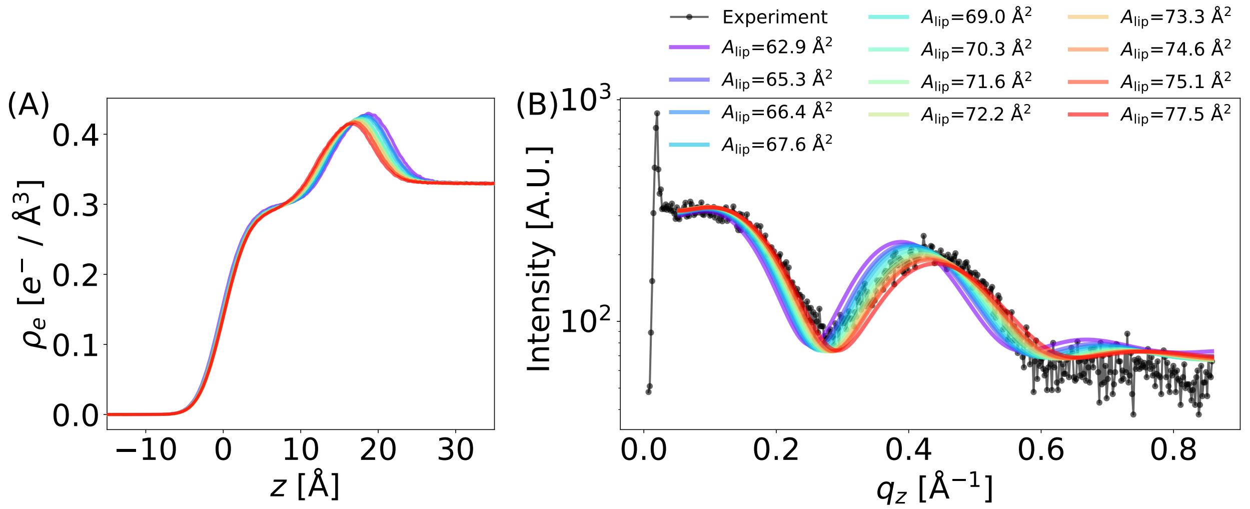


Figure S4: Comparison of simulated GIXOS curves at different values of $A_{\text{lip}}$ with experimental profile. Here the experimental profile correspond to pH 5.0.

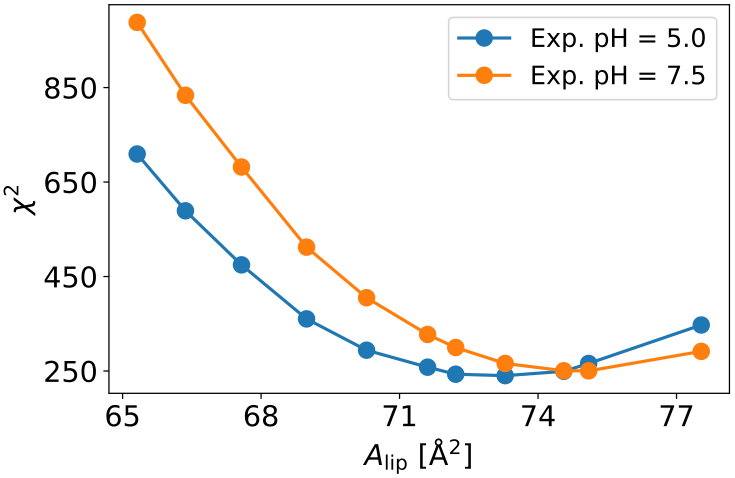


Figure S5: $\chi^2$ with varying area per lipid ($A_{\text{lip}}$) for pH 5.0 and pH 7.5. The optimum $A_{\text{lip}}$ corresponds the ones for which $\chi^2$ is minimum.

#### Comparison of GIXOS curves at 100 % $MC3^+$ and at correct protonation degree

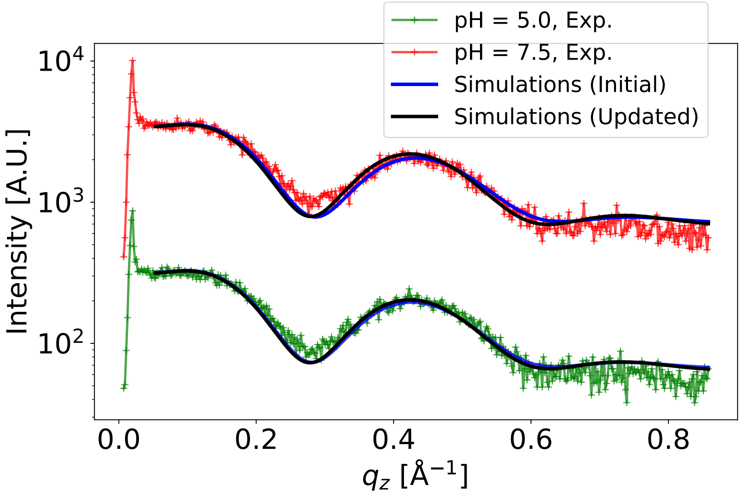


Figure S6: Comparison of simulation GIXOS curves at $\alpha$ = 100% ("Initial") and at the experimentally obtained protonation degree ("Updated").

#### Comparison of cation number density profile at at 100 % $MC3^+$ and at correct protonation degree

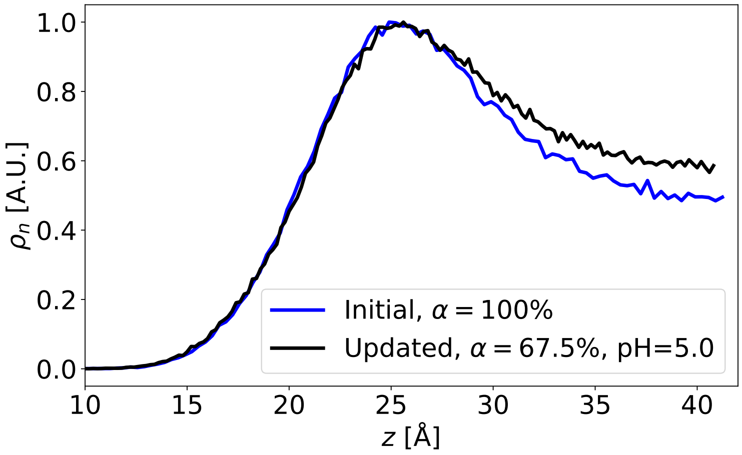


Figure S7: Comparison of cation profile with 100 % $MC3^+$ ("Initial") and at experimentally obtained protonation degree i.e, 67.5% $MC3^+$ ("Updated").

#### Density profiles of chemical components at protonation degree $\alpha = 15\%$

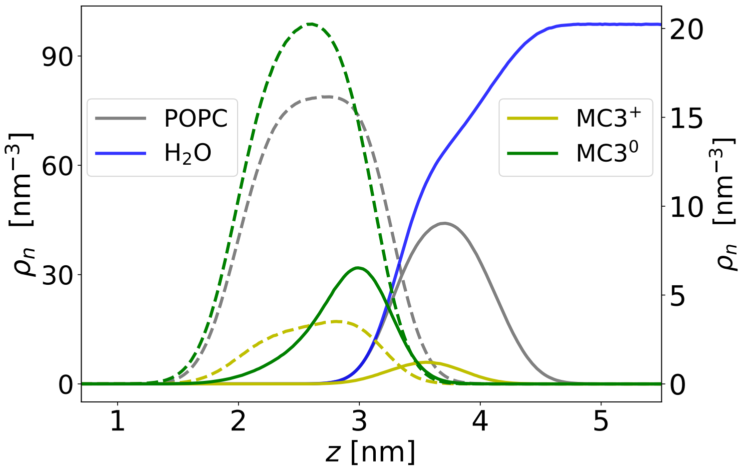


Figure S8: Number density of the chemical components at a protonation degree of $\alpha = 15\%$. The dashed curves represents the tail groups whereas full lines represents head groups. Here, the left vertical scale is for $H_2O$ and POPC and the right vertical scale is for heads and tails of $MC3^+$ and $MC3^0$.

#### 3D-radial distribution functions (RDFs)

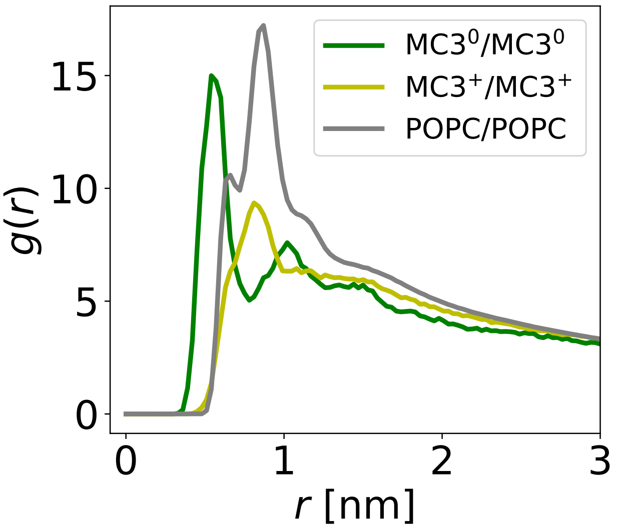


Figure S9: 3D-RDFs between the headgroup nitrogens of $MC3^0$, of $MC3^+$, and of POPC.

#### Potential of mean force for $MC3^0$ interactions

The potential of mean force (PMF) was obtained by Boltzmann inversion of the 2D-radial distribution function $g(r)$: $\mathrm{PMF}(r) = -k_{\mathrm{B}}T \ln(g(r))$. The PMF between $MC3^0$ head groups is shown in Figure S10 A. The PMF was smoothed by fitting a cubic spline function and the mean force was obtained by the negative gradient of the PMF (Figure S10 B). Figure S10 C shows a simulations snapshot with the repulsive and attractive force regions.

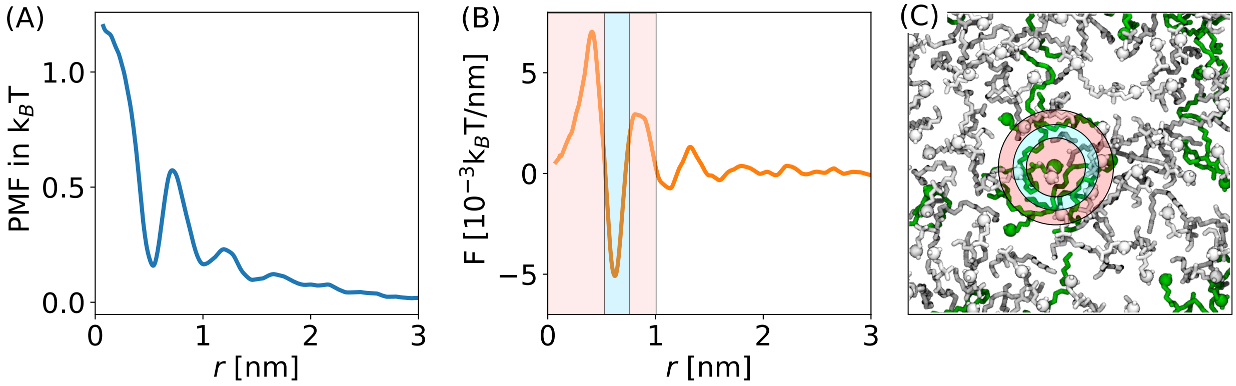


Figure S10: (A) Potential of mean force (PMF) between $MC3^0$ head groups. (B) Mean force between $MC3^0$ head groups exerted by all molecules within distance $r$. The shaded regions depict repulsion (light red) and attraction (light blue). Beyond $r > 1$ nm, the mean force vanishes. (C) Snapshot showing the repulsive and attractive force regions from (B). MC3 is shown in green and POPC in white. Head group nitrogens are depicted as spheres.
